# Off-season sex in *Zymoseptoria tritici*: little room for late encounters

**DOI:** 10.1101/2025.08.05.668626

**Authors:** Frédéric Suffert, Manon Delanoue, Stéphanie Le Prieur, Alicia Noly

**Affiliations:** Université Paris-Saclay, INRAE, UR BIOGER, 91120 Palaiseau, France

**Author notes:** Corresponding author: E-mail address (F. Suffert). The authors are listed in descending order of the importance of their contributions.

**Keywords:** avirulence, infection, plant disease epidemiology, Septoria tritici blotch, sexual reproduction, wheat residues

## Abstract

This study complements a substantial body of experimental work on the model fungal pathogen *Zymoseptoria tritici* by exploring marginal processes and conditions of sexual reproduction. Recent findings have shown that avirulent strains can engage in sexual reproduction on resistant host plants, even when they fail to cause visible symptoms during the biotrophic phase. The objective was here to examine the (de)coupling of various processes operating during the epidemic period (living plants) and the interepidemic period (crop residues). We assessed whether various encounter scenarios between parental strains, involving co-inoculations performed either simultaneously or sequentially on live and dead wheat plants, could result in successful mating. A two-year experiment accounted for the compatibility between wheat varieties (one carrying the resistance gene *Stb16q*, the other not) and the strains (virulent or avirulent), and the nature of the inoculum (blastospores or pycnidiospores). The intensity of sexual reproduction was assessed for each scenario through quantification of ascospore production, complemented by genotyping of offspring to confirm their parental origin. The main result is that a *Z. tritici* strain arriving late on dead host tissues can still mate with a compatible strain that previously colonized the plant, whereas sexual reproduction does not occur if both strains arrive after the plant has dried. Quantitative analysis suggests that although matings initiated by late encounters on wheat residues are possible, they contribute very little to the overall offspring population (< 2%). We discuss the epidemiological implications of this finding for disease management, highlighting both fundamental and applied questions it raises.

**Highlights:**

- Confirms avirulent *Z. tritici* strains reproduce sexually on resistant wheat.
- Late-arriving strains on dead tissues can mate with strains that infected earlier.
- Sex from such late encounters on wheat residues contributes little to offspring.
- Findings could impact understanding of pathogen lifecycle and disease control.

## 1. Introduction

Sexual reproduction in phytopathogenic ascomycete fungi remains enigmatic in many respects, both in terms of its physiological determinants and its epidemiological consequences. The physiological mechanisms occurring during sexual reproduction have been described in some model ascomycetes (e.g. *Neurospora crassa*; Brun *et al*., 2021), but the interaction with the host plant remains poorly understood for biotrophic and hemibiotrophic plant pathogens. Its impact is significant: qualitatively, it plays a key role in shaping population structure by facilitating genetic recombination and thus enhancing diversity (Ni *et al*., 2011); quantitatively, it contributes to the production of survival structures and primary inoculum, which are critical for the recurrence of epidemics (Burdon and Laine, 2019). Nevertheless, research on sexual reproduction remains neglected. In agronomy-oriented academic research, sexual reproduction is often avoided due to the experimental challenges it entails — particularly the need to fully understand the complex interactions between the pathogen and its host at specific developmental stages, which present significant methodological difficulties. Furthermore, sexual reproduction is rarely addressed in crop management strategies, as its effects on agricultural systems are indirect — or at least, seem to be so.

Septoria tritici blotch in wheat (STB) is caused by the heterothallic ascomycete fungus *Zymoseptoria tritici* (Suffert et al., 2011). Managing this major disease in Europe — where the deployment of resistant cultivars continues to face a steady erosion of effectiveness, despite ongoing improvements in their genetic background over the past three decades — remains a significant challenge (Fones & Gurr, 2015). The issue of sexual reproduction, as outlined just above, was framed for *Z. tritici* around two key points by Orellana-Torrejon et al. (2022) in the introduction of a study that served as a precursor to the present work. First, sex plays a crucial role in the multiannual recurrence of epidemics by generating variable amounts of STB inoculum on infected stubble residues, which are key to initiating and driving early-season epidemics (Morais et al., 2016; Suffert and Sache, 2011). In addition to this quantitative effect, sex has a qualitative impact on the evolutionary dynamics of the pathogen population — particularly through changes in population properties driven by the frequency and combination of alleles of interest, which are involved in selection processes that enable the pathogen to overcome host resistance genes and reduce the efficacy of fungicide treatments (McDonald & Mundt, 2016; Garnault et al., 2021; Kildea et al., 2021). Despite this, strategies targeting the sexual phase of the pathogen’s life cycle are largely absent from current STB management practices. Several studies have investigated the potential to act on the sexual reproduction phase to manage both inoculum levels and the evolutionary dynamics of the pathogen — through strategies such as reduced tillage (Kerdraon et al., 2019a), deploying cultivars in mixture (Orellana-Torrejon et al., 2022; Suffert et al., 2024), biocontrol based on microorganisms (Kerdraon et al., 2019b), but also components of the soil microfauna such as mycophagous springtails (Bourgeois et al., 2023). Nevertheless, what occurs during the interepidemic period — a key stage for fully leveraging agroecological approaches — has not received sufficient attention, and our overall understanding of the epidemiological processes taking place during this phase remains limited. In particular, little is known about the nature of interactions between the pathogen and the host plant — whether in terms of gene-for-gene virulence/resistance interactions or more complex dynamics — that might inhibit or facilitate sexual reproduction. These interactions also affect the transmission of certain alleles — such as those conferring virulence against major resistance genes — within pathogen populations, potentially accelerating the resistance erosion of the most commonly deployed wheat cultivars (Marliac et al., in prep).

Physiological and epidemiological processes are often studied separately — partly because this is more practical from a conceptual standpoint, and also due to limited biological knowledge linking the two, especially given that they often act on different spatiotemporal scales. This separation has its advantages: it allows for simplified modelling and helps to identify the influence of dominant processes that significantly shape the behavior of the pathosystem we aim to understand. However, this approach also has its drawbacks. By treating these processes independently, we risk overlooking so-called “minority” mechanisms — those that may be quantitatively marginal but could, in specific contexts, limit our ability to advance the understanding of interaction biology. This issue is particularly relevant to sexual reproduction in *Z. tritici*, which raises several basic yet important questions: which types of strain are involved, under what conditions, and with what consequences?

From an epidemiological standpoint — that is, considering the role a process plays during the course of an epidemic — sexual reproduction in *Z. tritici* is traditionally thought to occur solely on crop residues. Consequently, it is presumed not to interfere with the development of STB during the wheat growing season. Yet, this is now known to be either incorrect or, at best, an oversimplification, as sexual reproduction has also been observed on the lower leaves of living plants (Eriksen & Munk, 2003; Suffert & Sache, 2011). What are the determinants of that, and what are the implications? To date, no definitive answers exist. From a physiological standpoint, sexual reproduction in *Z. tritici* is typically considered a secondary phase, even at the scale of individual wheat leaf tissues, and is assumed to occur only after asexual reproduction has taken place. This assumption is based on the view that only the asexual infection phase — particularly in a hemibiotrophic pathosystem such as that of STB — allows for the coexistence of strains, which may then lead to sexual reproduction if they are compatible, i.e., of opposite mating types. However, recent research challenges this dual view. Various crossing scenarios have also explored how the nature of compatible or incompatible interactions during the asexual phase may influence the outcome of sexual reproduction (Kema et al., 2018; Orellana-Torrejon et al., 2022) and it is now known that the symptomatic, asexual infection phase is not a prerequisite for at least one of the two parental strains involved in sexual reproduction. Avirulent strains can engage in sexual reproduction on resistant host plants, even when they do not cause any visible symptoms during the biotrophic phase — in other words, two avirulent strains can mate. Altogether, these empirical findings reshapes the population genetics baseline of this fungus — and likely that of many other Dothideomycetes. This idea deserves to be more widely integrated by the phytopathology community, even though it represents a shift in established dogma.

The objective of this experimental study was to deepen our understanding of sexual reproduction processes in *Z. tritici*, with a particular focus on the (de)coupling of mechanisms operating at the plant scale between the epidemic period (on living plants) and the interepidemic period (on crop residues, where primary inoculum is formed as a result of sexual reproduction). We aimed to determine whether two *Z. tritici* strains are capable of mating following late contact on wheat residues — i.e., on dead plant tissue — rather than during earlier interactions on living plants. While the question is simply stated and may appear straightforward, it raises both experimental and epistemological challenges. On one hand, detecting processes that occur at very low frequencies requires careful management of “false positives”: how can we be confident that a rare event is genuine and not the result of an artefact or misinterpretation? On the other hand, establishing that a process does not occur is equally problematic due to the risk of “false negatives”: it is virtually impossible to formally prove non-existence — it may simply occur below the detection threshold of the experimental design.

## 2. Materials & methods

### 2.1. Overall strategy

The experimental strategy was designed to test whether two *Z. tritici* strains can mate when they come into late contact on wheat tissues that are dead and thus considered to be residues. A crossing plan was implemented to evaluate this scenario by comparing it with others, taking into account three key determinants — either known or suspected — of sexual reproduction in STB (**Figure 1**). These factors included: (i) sexual compatibility of the parental strains, (ii) compatibility between the host and pathogen (gene-for-gene interaction), and (iii) the form in which the pathogenic inoculum is introduced upon contact with the plant (blastospores vs. pycnidiospores). Pairs of virulent and/or avirulent strains with opposite mating types were used. Inoculations were carried out on two wheat varieties — one with a major resistance gene and one without — either in a staggered manner over time or simultaneously. In the staggered inoculation scenario, a single strain was initially introduced to a living adult plant, which either developed symptoms or remained asymptomatic depending on the virulence status of this strain relative to the cultivar; the second strain was then introduced at a later stage, once the plant had dried out (scenario PR). In the simultaneous inoculation scenario, both strains were inoculated at the same time, either onto a living adult plant (scenario PP) or onto a plant that had been left to dried out and considered as impending residues (scenario RR). For all crossing scenarios, the fully dried plants were prepared as bundles of residue sets and placed outdoors for five months to promote ascosporogenesis. The intensity of sexual reproduction was estimated by quantifying ascospore discharge (Suffert et al., 2016). Several sexually derived offspring strains were recovered from selected residue sets and genotyped using SSR markers to verify their ‘parenthood’ (Gautier et al., 2014). Their genotypic profiles were compared with those of the co-inoculated strains to identify which ones had actually undergone crossing (Orellana-Torrejon et al., 2022).

**Figure 1.**
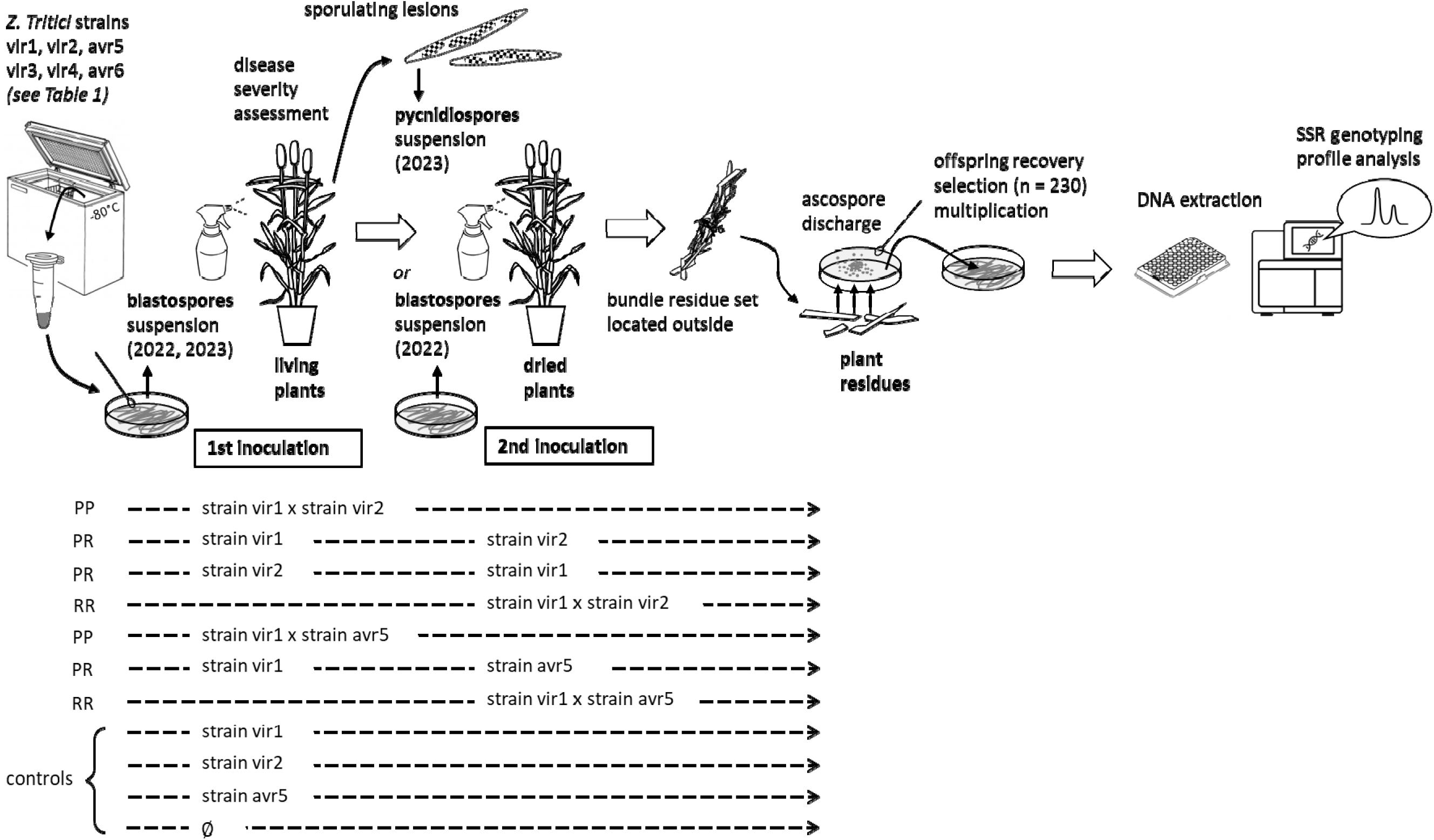
Experimental strategy and standard crossing scheme.

The experiment was conducted in the first year (2022) using blastopores — asexual spores produced by budding in yeasts — produced *in vitro* for all inoculations. In the second year (2023), initial inoculations of living adult plants were performed using blastospores, whereas those of dried plants were later conducted with pycnidiospores — asexual spores released in cirrhi extruding from pycnidia — collected from sporulating lesions on wheat plants that had been specifically prepared for this purpose. In both years, two parallel experimental series were conducted as biological replicates, each on a separate greenhouse compartment. Each series involved a different trio of parental strains with similar virulence statuses: vir1, vir2, and avr5 in one; vir3, vir4, and avr6 in the other (**Figure 1**).

### 2.2. Plant and fungal material

We used the wheat cv. Cellule (Florimond Desprez, France), which carries the *Stb16q* resistance gene, and the susceptible cv. Apache (Limagrain Europe, France). Two-week-old seedlings were vernalized in a growth chamber at 8°C for 8 weeks, then transplanted into individual pots and grown under the same conditions described by Orellana-Torrejon et al. (2022). Plants were thinned to three stems per pot during the growth period.

Six *Z. tritici* parental strains were selected (**Table 1**), all collected from a wheat field in Grignon, France, in 2018. Their sexual compatibility was determined through PCR amplification of the two mating-type idiomorphs (Waalwijk et al., 2002), while their virulence status concerning *Stb16q* was assessed by *in planta* phenotyping (Orellana-Torrejon et al., 2022). Blastospore suspensions of each strain, preserved at-80°C, were prepared from subcultures grown for five days on Petri dishes containing PDA (potato dextrose agar; 39 g·L^−1^) at 18°C in the dark, following the protocol of Suffert et al. (2013). The pycnidiospore suspensions used in the second year to inoculate dried plants were prepared by suspending in water leaf fragments with sporulating lesions, which were collected from plants six weeks after they had been specifically inoculated as previously described (**Figure 2A**). All inoculum concentrations were adjusted to 5 × 10L spores·mLL¹ using a Malassez counting chamber, and two drops of surfactant (Tween 20; Sigma, France) were added. Two-strain suspensions, consisting of a Mat1.1 strain and a Mat1.2 strain, were prepared by mixing equal volumes (1:1) of the corresponding single-strain suspensions.

**Figure 2.**
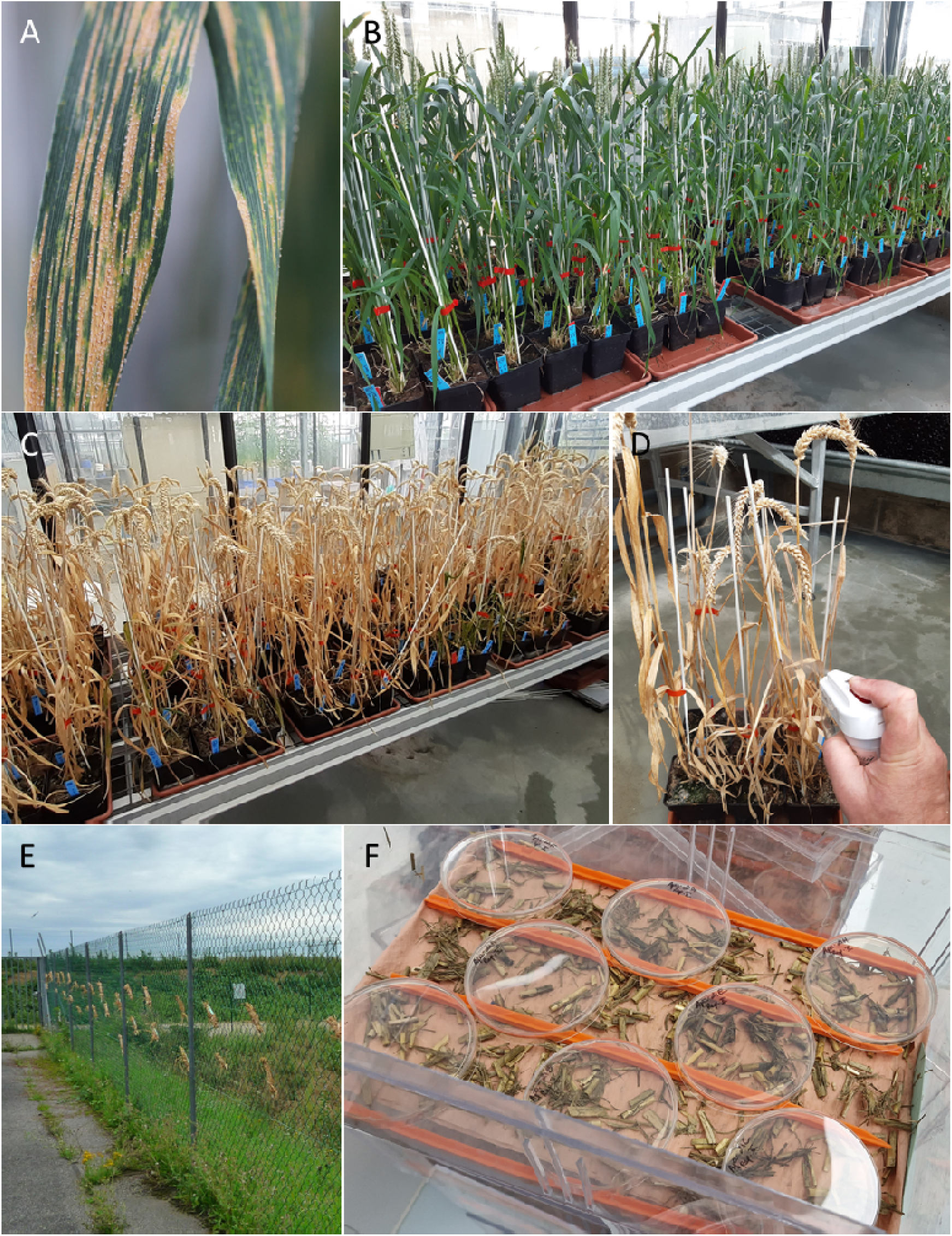
**A.** Septoria tritici blotch lesions. **B.** Living wheat plants (cv. Cellule and Apache) before the first inoculation. **C.** Dried wheat plants before the second inoculation. **D.** Inoculation of dried plants using an atomizer. **E.** Bundles of residue sets hung on a fence in autumn to promote ascosporogenesis. **F.** Petri dishes containing PDA medium placed upside down above pieces of wheat residue to collect *Zymospetoria tritici* ascospores.

**Table 1.** Parental strains of *Zymoseptoria tritici* used in crosses.

| Replicate | Id | Strain | Mating type | *Stb16q* virulence status |
| --- | --- | --- | --- | --- |
| rep1 | vir1 | INRA18-FS3250 | MAT1.1 | virulent |
|  | vir2 | INRA18-FS2883 | MAT1.2 | virulent |
|  | vir3 | INRA18-FS3098 | MAT1.2 | virulent |
| rep2 | vir4 | INRA18-FS2881 | MAT1.1 | virulent |
|  | avr5 | INRA18-FS2849 | MAT1.2 | avirulent |
|  | avr6 | INRA18-FS2851 | MAT1.1 | avirulent |

### 2.3. Inoculation procedure

An atomizer (Ecospray, VWR, France) was used to apply each inoculum suspension of pycnidiospores onto three adult plants (nine stems) of the cv. Apache and Cellule after the wheat heads had fully emerged (on 19^th^ April 2022 and 30^th^ May 2023; **Figure 2B**), as described by Suffert et al. (2013) and following the experimental design presented in **Figure 1**. The second inoculations on dried plants were performed once the plants had reached full senescence (**Figure 2C**), also in the greenhouse, with suspensions of blastospores (on 1^st^ August 2022) or suspensions of pycnidiospores (on 7^th^ August 2023). In all cases, 20 mL of inoculum suspension was sprayed onto each trio of plants while rotating them for approximately 30 seconds to ensure even coverage with inoculum (**Figure 2D**). Infection was promoted by immediately enclosing the plants, whether still living or dried, in a transparent polyethylene bag containing a small amount of distilled water for 72 hours after inoculation. To avoid cross contamination the pots of the different trio of plants were then separated by about 20 cm to prevent their leaves from touching. Air temperature was maintained above 25°C until June, with occasional peaks reaching 35°C in July and August. Adult plants of both cv. Apache and Cellule were used as negative (non-inoculated) controls.

We estimated the intensity of asexual reproduction by assessing STB severity — defined as the percentage of leaf area covered by pycnidia (1%, 2%, 3%, 5%, and in 5% increments thereafter up to 100%; Suffert et al., 2013) — on the two uppermost leaves of each stem of the inoculated plants, five weeks after inoculation (on 21^th^ May 2022 and 3^th^ July 2023). The pathogenicity of the strains — their virulence or avirulence status on the two cv. Apache and Cellule — was verified through a qualitative assessment of disease severity, based on the presence or absence of symptoms. A quantitative analysis was performed using ANOVA to assess the effects of inoculum type (confounded with year), strain combinations, and cultivar on the severity, complemented by a non-parametric Mann–Whitney U test.

### 2.4. Promotion and assessment of sexual reproduction

Three days after the final inoculation of dried plants (on 4^th^ August 2022 and 10^th^ August 2023), each trio of plants was bundled into small packages, as described by Orellana-Torrejon et al. (2022), and hung on a wire mesh outdoors during the summer and autumn to promote ascosporogenesis (**Figure 2E**). After five months of exposure to outdoor weather conditions, the senescent leaves and stems from each bundle residue set were collected (on 8th January 2022 and 19^th^ December 2023), cut into 2 cm segments, and left to dried at 18 °C in laboratory conditions for some days.

The intensity of sexual reproduction on each set of residues was assessed by estimating ascospore discharge events, with each event repeated once. Residues were weighed, soaked in water for 30 min and spread on dried filter paper in a box (24 × 36 cm), the lid of which was left half-open to gradually decrease the relative humidity. Eight Petri dishes (90 mm in diameter) containing PDA medium were opened and placed upside down 1 cm above the residues at 19°C for 24 h (**Figure 2F**). The dishes were then closed and incubated in the dark at 18°C for five days. The number of ascospore-derived colonies was estimated visually to calculate a cumulative ascospore discharge index (ADI; ascospores·glJ¹), defined as the total number of ascospores discharged per gram of wheat residues, summed over the two discharge events (Suffert & Sache, 2011; Suffert et al. 2016; Orellana-Torrejon et al., 2022).

### 2.5. Offspring genotyping

A total of 229 offspring strains were sampled in the Petri plates from various ascospore-derived colonies obtained from successful crosses conducted during the 2022 and 2023 experiments. The goal was to capture the diversity of crossing scenarios, with particular emphasis on those involving inoculations on dried plant (notably scenarios PR), while limiting the sampling to a maximum of 15 offspring per cross. The strains were subcultured onto PDA and stored in cryotubes at-80°C. Their parenthood was determined by comparing their genotyping profiles with those of the strains used for (co-)inoculation and presumed to be their parents (**Supplementary Table S1**). Twelve SSR markers (AC0002-ST4, AG0003-ST3A, CT0004-ST9, TCC0009-ST6, AC0001-ST7, GCA0003-ST3C, chr_02-140-ST2, GAC0002-ST1, GGC0001-ST5, CAA0003-ST10, TCC0002-ST12, CAA0005-ST3B) from the panel developed by Gautier *et al*., (2014) were used. DNA extraction was performed on 5-day-old strain cultures using a 10% Chelex solution for the 2022 offspring and using the Qiagen DNeasy Plant Maxi Kit for the 2023 offspring. Allele identification for each marker was based on sequencing results of each extract, which were conducted by Eurofins SA, followed by verification using Peak Scanner software v2.0 (Applied Biosystems). The profile comparison was carried out using the decision tree developed by Orellana-Torrejon et al. (2022). An offspring was definitively considered not to be the result of the cross between the two co-inoculated strains if at least one of its 12 SSR markers did not match those of either strain. Conversely, if all 12 markers matched, it was deemed highly probable that the offspring originated from the directed cross. For each crossing scenario, the degree of exogeneity of the actual parental strains was estimated by the percentage of genotyped offspring samples that have at least one parent different from the co-inoculated strains.

### 2.6. Analysis of the sexual reproduction intensity across different crossing scenarios

The analysis aimed to identify the scenarios that led to *Z. tritici* crosses, with a particular focus on those involving the inoculation of dried wheat tissues considered to be at the residue stage. To enable a quantitative comparison, we evaluated the intensity of sexual reproduction, estimated using the cumulative ADI proxy, for each crossing scenario — simultaneous co-inoculations on living plants (PP), co-inoculations on dried plants (RR), and separate inoculations on each plant type (PR) — accounting for their respective weighting in the experimental design (twice as many PR as PP and RR). We distinguished between the types of inoculum applied to dried plants (blastospores in 2022 vs. pycnidiospores in 2023). A preliminary quantitative analysis was carried out using multivariate ANOVA to assess the effects of replicate (set of strains), inoculum type (confounded with year), wheat tissue type, and crossing scenario type. Since the data did not follow a normal distribution (according to the Shapiro-Wilk test) and exhibited heterogeneous variance (based on Levene’s test), the impact of crossing conditions — whether characterized by the wheat tissues type or the crossing scenario type — on the intensity of sexual reproduction (ADI) was analysed using a non-parametric Mann-Whitney test. The overall results were discussed in light of the genotyping data, which provided strong indications regarding the origin of the offspring resulting from the crosses.

By extrapolating the results, we ultimately estimated the potential origin of crosses occurring in plant residues remaining on the soil surface after wheat harvest, under the assumption that inoculum is not limiting — neither during the growing season (pycnidiospores produced from living wheat plants) nor during the interseason (pycnidiospores produced from crop residues on the ground) — and that both inoculum sources contribute equally.

## 3. Results

### 3.1. STB severity levels analysed as indicators of the nature of the strain-cultivar interaction during the asexual phase

The virulence status of the six *Z. tritici* parental strains with respect to the two cultivars was verified through a qualitative assessment of STB severity (presence or absence of symptoms) following single-strain inoculations used as control. All strains caused symptoms on cv. Apache, whereas only the four virulent strains produced symptoms on cv. Cellule. The impact of virulence status of strains single-or co-inoculated was also quantitatively evident, as shown by differences in STB severity between Cellule and Apache following inoculation while adult plants were still alive. ANOVA (**Table 2**) revealed significant effects of inoculum type (confounded with year) and cultivar — with Apache appearing more susceptible. The main effect of strain combination was marginally significant (p ≈ 0.0603). A highly significant interaction between parental strain combinations and cultivar was observed: on Cellule, strain combinations had a strong influence on STB severity, whereas on Apache this effect was much weaker or even absent. These results should be interpreted with caution, as the assumption of residual normality was met (Shapiro-Wilk test; p = 0.074), but the assumption of homogeneity of variances was not (Levene’s test; p = 0.0083).

**Table 2.** Results of the multivariate ANOVA aimed at exploring the effects of inoculum type (blastospores in 2022, pycnidiospores in 2023), cultivar (Apache, Cellule), strain combination, and significant interactions, on STB severity.

| Factor | Sum of squares (SS) | ddl | F | p-value |
| --- | --- | --- | --- | --- |
| Inoculum type | 1858.72 | 1 | 9.64 | 0.0034 |
| Cultivar | 9248.70 | 1 | 47.96 | <0.0001 |
| Strain combinaison | 3503.93 | 9 | 2.02 | 0.0603 |
| Cultivar × strain combinaison | 11752.70 | 9 | 6.77 | <0.0001 |
| Residues | 8292.35 | 43 | — | — |

The Mann–Whitney U test comparing the STB severity obtained for each strain combination between Apache and Cellule detected significant differences only for single inoculations with avr5 and avr6 (p = 0.029), and none for the co-inoculations. Visually, a large difference — close to a 50% magnitude — was observed for the combinations vir1 × avr5 and vir3 × avr6 (**Figure S1**), but it was not statistically significant due to the small sample size. The overall comparison of STB severity between Apache (54.2 %) and Cellule (30.2 %) was highly significant (Mann–Whitney test; p < 0.001). For single-or co-inoculations with only virulent strains, the difference in STB severity observed on Apache and Cellule was not significant. This corresponds to the expected behavior of the strains used.

### 3.2. Analysis of crossing scenarios that led to mating, following confirmation by genotyping of the offspring

The distribution of the ADI variable, which reflects the intensity of sexual reproduction, was not normal (Shapiro-Wilk test; F = 0.412; p < 0.001), and the hypothesis of homogeneity of variances was rejected, whether focusing on the effect of the wheat tissues type (Levene’s test; F = 24.49; p < 0.001) or the effect of crossing scenario type (Levene’s test; F = 13.52; p < 0.001). This justifies the use of the Kruskal-Wallis test, the results of which show a significant effect of both factors studied (F = 32.35 and F = 33.43, respectively; p < 0.001). A multivariate ANOVA, conducted as a complementary analysis, reveals complex interactions and significant effects of inoculum type, as well as highly significant effects of crossing conditions, whether characterized by the type of wheat tissues inoculated or the crossing scenario (**Table 3**).

**Table 3.**
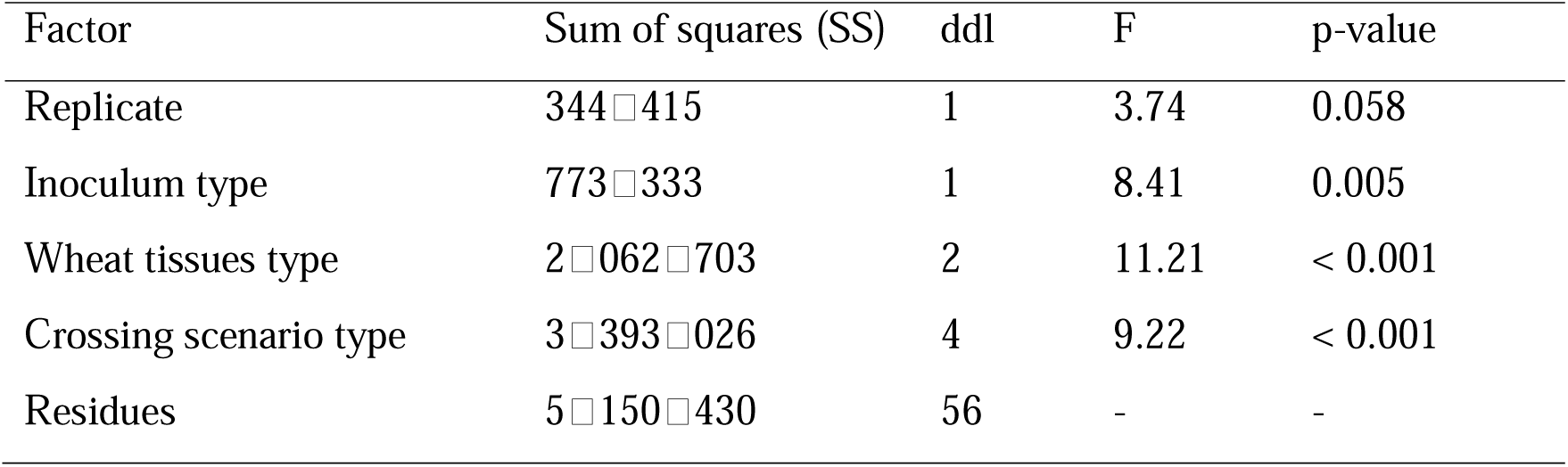
Results of the multivariate ANOVA aimed at exploring the effects of replicate (set of strains), inoculum type, wheat tissues type, and crossing scenario type on the intensity of sexual reproduction, estimated by the cumulative ascospore discharge index (ADI).

| Factor | Sum of squares (SS) | ddl | F | p-value |
| --- | --- | --- | --- | --- |
| Replicate | 344□415 | 1 | 3.74 | 0.058 |
| Inoculum type | 773□333 | 1 | 8.41 | 0.005 |
| Wheat tissues type | 2□062□703 | 2 | 11.21 | < 0.001 |
| Crossing scenario type | 3□393□026 | 4 | 9.22 | < 0.001 |
| Residues | 5□150□430 | 56 | - | - |

Co-inoculations performed simultaneously on living adult plants with blastospores (PP scenario) successfully produced offspring in nearly all cases 15 out of 16 in both 2022 and 2023. The average recovery was 575 ascospores per gram of residue (cumulative across the two discharge events), with a notable difference between the two years: 121 ascospores per gram in 2022 versus 956 ascospores per gram in 2023. Among these, all the expected crosses (8 out of 8 cases) — those in which the co-inoculated strains were both sexually compatible and virulent with respect to the host plant cultivar (vir × vir on Cellule and vir × avr on Apache) — were successful (**Supplementary Tables S1 and S2**). Analysis of genotyping data from offspring sampled from these 8 crosses confirmed, with high confidence, that the parental strains involved in sexual reproduction were the co-inoculated strains, with no evidence of contribution from exogenous ones.

Scenarios involving co-inoculation with one virulent and one avirulent strain with respect to the inoculated cultivar (3 out of 4 cases) still resulted in successful crosses. A strong effect of the inoculum type, confounded with a year effect, was observed: in 2022, no successful crosses were recorded, except for the vir1 × avr5 combination on Cellule, which produced very few ascospores (1.1 ascospores per gram of residue), and only three offspring could be genotyped. In contrast, in 2023, the vir1 × avr5 and vir3 × avr6 crosses on Cellule were successful, yielding a large number of offspring. Genotyping of 15 and 16 individuals, respectively, confirmed that the parental strains involved in sexual reproduction were very likely the co-inoculated strains.

Scenarios involving an initial inoculation of the first strain on living adult plants followed by a delayed second strain inoculation when the plants were dried (PR) also resulted in successful crosses: only 1 out of 16 cases in 2022, but 11 out of 16 in 2023. Offspring samples were analysed for nine of these crosses, and in five cases, the strain inoculated second was very likely one of the two parents. For the remaining four crosses, certainty is lower, as some offspring alleles did not completely match those of the inoculated strains, suggesting that at least one parent originated from an exogenous origin. The higher number of successful crosses in PR scenarios in 2023 — corresponding to the use of pycnidiospores — is supported by the ANOVA results, but this finding should be interpreted with caution, given that the assumptions underlying this type of analysis were not fully met. Co-inoculations performed on dried plants (RR scenario) did not yield any offspring, except in 2023, when three individuals were collected from a single vir3 × vir4 cross on Apache, corresponding to 0.3 ascospores per gram of residue (**Supplementary Table S1**). Genotyping of the offspring was not performed, as the results — regardless of the outcome — would have been inconclusive given the very small sample size.

Finally, inoculations with single-strain suspensions resulted in offspring in 6 out of 16 cases, with the exogenous origin of one of the two parents being unequivocal, as confirmed by the genotypes of the offspring obtained in two of these crosses (**Supplementary Table S1**). Although this proportion appears high, the number of ascospores recovered in each case was low, representing less than 6% of the total offspring.

### 3.3. Quantitative analysis of the impact of crossing scenarios on sexually derived offspring

ADI was significantly influenced (Mann-Whitney U test) by the crossing conditions — whether characterized by the wheat tissues type at the time of each inoculation or the crossing scenario type following the categorisation proposed by Orellana-Torrejon et al. (2022). Regardless of the inoculum type used for the second inoculation, 98.6% of ascospores were collected in PP scenarios, 1.4% in PR scenarios, and none in RR scenarios (**Table 4**). The distribution by crossing scenario shows that the vast majority of ascospores were recovered in types 1 and 2 (approximately 98% combined). Although the proportion between types 1 and 2 varied greatly between 2022 and 2023, the differences were not statistically significant due to the low number of crosses associated with type 2. Around 1% of ascospores were recovered in types 3 and 4, and none in type 5.

**Table 4.** Mean intensity of sexual reproduction estimated by the cumulative ascospore discharge index (ADI), according to crossing conditions characterized by (**A**) the wheat tissues type at the time of each parental strain inoculation (P = living plants; R = dried plants) and (**B**) the crossing scenario type (1 to 5) following the categorisation proposed by Orellana-Torrejon et al. (2022), taking into account the nature of the inoculum (blastospores or pycnidiospores). The data from both replicates (rep1 and rep2) were merged, and detailed results are available in **Supplementary Table S1**. Different letters indicate that all crossing conditions differ significantly from each other, according to multiple Mann Whitney U tests with Bonferroni correction (p < 0.01).

| Crossing conditions |  | Inoculum type used for the second (co)-inoculation, if applicable <sup>1</sup> |  |  |  |
| --- | --- | --- | --- | --- | --- |
|  |  | Blastospores (2022) |  | Pycnidiospores (2023) |  |
| (A) Wheat tissues type | PP | 121.3 (98.6 %) | a | 956.2 (98.6 %) | a |
|  | PR | 1.8 (1.4 %) | b | 13.9 (1.4 %) | b |
|  | RR | 0.0 (0.0 %) c | b | 0.0 (0.0 %) | c |
| (B) Crossing scenario type <sup>2</sup> | 1 | 161.6 (97.9 %) | a | 864.2 (40.7%) | a |
|  | 2 | 0.6 (0.3 %) | a | 1232.2 (58.0 %) | a |
|  | 3 | 2.0 (1.2 %) | b | 14.0 (0.7 %) | b |
|  | 4 | 0.0 (0.0 %) | b | 13.4 (0.6 %) | - |
|  | 5 | 0.0 (0.0 %) | b | 0.0 (0.0 %) | c |

<sup>1</sup> wheat tissues type PR and RR; crossing scenario types 3, 4 and 5.
<sup>2</sup> 1 = both parental strains are virulent and can infect independently living wheat tissues; 2 = one parental strain is virulent and can infect living wheat tissues, whereas the other parental strain can only grow asymptotically, possibly epiphytically, on living wheat tissues; 3 = one parental strain is virulent and can infect living wheat tissues, whereas the other parental strain (virulent or avirulent) arrives later on dried wheat tissues considered residues; 4 = one parental strain is avirulent and can only grow asymptotically, possibly epiphytically, on living wheat tissues, whereas the other parental strain (virulent or avirulent) arrives later on dried wheat tissues considered residues; 5 = both parents (virulent or avirulent strain) arrive late on dried wheat tissues considered residues.

## 4. Discussion

### 4.1. Strains that arrive late on dead tissue can still engage in sexual reproduction, but the number of offspring produced under these conditions is low

The main result, which addresses the objective of the experiment, is that a *Z. tritici* strain arriving on dry host tissues — tissues it is unable to infect — can still mate with a sexually compatible partner that has infected those tissues, thereby producing sexual offspring. This pattern was infrequent in 2022 (1 out of 16 cases) but more frequent in 2023 (13 out of 16 cases). Our results clearly show that successful mating under these conditions requires the other parental strain to have previously infected the host tissues, as indicated by the presence of STB symptoms. This is illustrated by the absence of offspring in the PR treatments when the first single inoculation involved an avirulent strain, and in the RR treatments. Only two observations could challenge this requirement: the production of some offspring from the avr6 × vir3 cross in the PR treatment in 2023, although parental origin is uncertain in this case due to an estimated 38% exogeneity; the recovery of offspring following co-inoculation with vir3 and vir4 on dry plants — but in an extremely low number (n = 3). Given the uncertainties associated with these exceptions, they should be interpreted with caution. The failure of co-inoculation on completely dry plants (RR) is consistent with the results of Halama & Lacoste (1992), who observed successful crosses of *Phaeosphaeria nodorum* on dry wheat straw, but not of *Z. tritici*. It also aligns with earlier negative results from greenhouse trials conducted at INRAE BIOGER, where rainwater was sprayed on dry plants for two months before they were moved outdoors (Suffert & Delanoue, unpublished data).

### 4.2. Results are supported by the behavior of avirulent strains and a robust methodology

The results are consistent with previous studies using the same methodology and inoculations performed exclusively on living adult plants. All crosses resulting from simultaneous co-inoculations on living adult plants were successful, including those where one of the two strains was avirulent on the wheat cultivar. Offspring were recovered, and genotyping of a subset confirmed, in the majority of cases, that they resulted from recombination between the two inoculated parental strains. The success of such crosses is in line with the findings of Orellana et al. (2022) and Kema et al. (2018). Such external validation was essential for drawing robust conclusions (regarding success or failure of sexual reproduction) in the context of inoculations on dry plants. The appearance of a few late-developing lesions following inoculation with avirulent strains (e.g., avr4, avr5, and avr6 on cultivar Cellule in 2023) illustrates that, as leaves begin to senesce, physiological barriers to infection may weaken, allowing the formation of some sporulating lesions — an observation also reported by Orellana et al. (2022). The single inoculations served to verify the virulence status (virulent vs. avirulent) of the six *Z. tritici* strains and to confirm the absence of contamination at the time of inoculation. That crosses also occurred — albeit at a very low level — in some of these single inoculation conditions may seem surprising at first. However, this too is consistent with findings by Orellana et al. (2022), who explained such events as the result of external strains arriving later, after the tissue of residue sets had been placed outdoors and had dried. The present study provides further evidence that directly supports this hypothesis (see below).

Genotyping of several series of offspring (around 15 per analysed cross) allowed verification of the parental strain origins. The simplified procedure for recovering offspring from isolated ascospore-derived colonies (involving only one intermediate single-spore subculturing) repeatedly led to the detection of double alleles among the tested offspring due to the presence of mixed-strain samples. This practice could be improved; however, it did not pose an issue in this case, as both alleles — when present — matched those of the inoculated strains.

### 4.3. Environmental conditions have a quantitative effect on the intensity of sexual reproduction

The difference in the intensity of sexual reproduction between the two years of the experiment was highly significant for the PP scenarios, with a 1-to-9 ratio in ADI: 121 ascospores per gram of residue following the second inoculation with blastospores, compared to 956 with pycnidiospores. One might be tempted to attribute this difference to the type of inoculum, but its effect on the intensity of sexual reproduction is confounded with a year effect due to the experimental design. Since the nature of the inoculation in the PP treatments was identical in both years (inoculation of living plants with blastospores), the observed difference in sexual reproduction intensity between 2022 and 2023 is most likely due to climatic conditions that the residue bundles were exposed to. In particular, this difference may be linked to cumulative rainfall in September and October (94 mm in 2022 vs. 142 mm in 2023), which is known to promote ascosporogenesis. The impact of rainfall on sexual reproduction had already been observed in a field trial conducted in Grignon, where the number of ascospores trapped in autumn 2012 was ten times higher than in 2011, corresponding to cumulative rainfall of 96 mm in 2012 versus 57 mm in 2011 (Morais et al., 2016). Since the ADI ratio between 2022 and 2023 for the PR treatments was of the same magnitude (1-to-9, with 2 ascospores per gram of residue versus 14), no definitive conclusion can be drawn regarding the effect of the inoculum type used in the second inoculation of dry plants (blastospores in 2022, pycnidiospores in 2023) on the intensity of sexual reproduction.

### 4.4. The physiological mechanisms underlying sexual reproduction remain unresolved

Our results show that crosses with a parental strain arriving late are only possible if the first parental strain has previously succeeded in infecting the tissues — achieved here through inoculation of living plants — as evidenced by the absence of offspring in the RR treatments. This result raises questions about the physiological mechanisms underlying sexual reproduction in *Z. tritici*. Some hypotheses have been proposed by Orellana et al. (2022), particularly regarding the diversity of configurations that can lead to mating. We do not revisit all the discussion points from that study, but it is worth recalling the hypothesis that, similar to *Magnaporthe oryzae* (Lassagne et al., 2022), microconidia may be involved at a late stage and might be preferentially produced alongside either type of spore (blastospore or pycnidiospore). This hypothesis could not be validated here, as sexual reproduction occurred in the PR scenario regardless of whether the second strain was introduced as blastospores (in 2022) or pycnidiospores (in 2023), and the differences in the intensity of sexual reproduction might be explained by a year effect. It is possible that, in each inoculum suspension, some small spores may have acted as male gametes. Significant gaps remain in our understanding of the interactions between living plants and pathogenic fungi during sexual reproduction, due in part to the experimental challenges in inducing sexual reproduction in some organisms and their ability to switch lifestyles (epiphytic, endophytic, and pathogenic; e.g., Schulz and Boyle, 2005). The inability to perform in vitro crosses, as has been done with *M. oryzae* (Saleh et al., 2012a, 2012b) or *Leptosphaeria maculans* (Shoemaker and Brun, 2011), severely limits our understanding of the biological mechanisms involved in sexual reproduction in *Z. tritici*. This is compounded by the difficulty of conducting cytological studies on dead plant tissues. Indirect approaches such as those used in this study — while inherently lengthy and complex — remain necessary, even though the advances in knowledge they produce may seem limited compared to the experimental efforts involved.

### 4.5. Estimating the relative likelihood of the different crossing scenarios under field conditions and their epidemiological consequences is possible

As illustrated in the title of this article, matings initiated by late encounters on wheat residues are possible but contribute very little to the offspring population. The quantitative analysis of our experimental results indeed reveals that the proportion of offspring obtained from the PR treatments is very low (<2%) compared to the PP treatments. Extrapolating from this, all other factors being equal, suggests that at the population level, no more than 2% of ascospores present in the air during autumn are produced by late-stage crossings. Such an extrapolation to field conditions is not straightforward, as the experiment involved an artificially balanced inoculum. In natural conditions, the number of pycnidiospores reaching wheat plants versus wheat stubble and residues — assuming these remain on the soil surface — may differ considerably. This difference stems partly from the distinct sources of inoculum, the varying amounts of receptive tissue (living plants vs. residues), and the fact that the efficiency of spore dispersal onto a substrate depends on environmental conditions. Therefore, the 2% figure likely overestimates the actual contribution in the field. In conclusion, while we have demonstrated that crossings occurring late on residues are possible, their epidemiological significance is most likely negligible.

The division into treatments PP, PR, and RR — based on the type of wheat tissue and the timing of co-inoculations (simultaneous or staggered) — was well suited to controlled conditions. In contrast, the division into five scenarios (1 to 5), four of which were described by Orellana et al. (2022), offers greater insight into what might occur under field conditions. The findings reported here are consistent with the results of that earlier study.

Scenarios of type 1 and 2 are two subtypes of the PP scenario, distinguished by whether both parents or only one are virulent. Type 1, in which both parents (virulent strains) are capable of independently infecting living tissue, is the most common and typically leads to sexual offspring. Type 2 involves one parent (a virulent strain acting as the maternal parent) infecting living tissue, while the other (an avirulent strain acting as the paternal parent) grows asymptomatically. This particular type of cross was specifically studied by Orellana et al. (2022), who demonstrated that such cross is possible and plays a non-negligible role under certain cultivation conditions, such as varietal mixtures.

Scenario of type 3 corresponds to the PR treatment, with the first parent (a virulent strain acting as the maternal parent) that infects the wheat plant while it is still alive. This type may explain cases where single inoculations led to crossing, and is also comparable to the PP scenario when one of the two parents is avirulent. In this context, “one parent (virulent strain) infects living tissues, whereas the other parent (virulent or avirulent strain) arrives later on crop residues — either via pycnidiospores from nearby residues or via ascospores traveling longer distances — and then grows saprophytically” (Orellana et al., 2022). Type 3, although shown in this study to be rare, can occur during the interepidemic period in autumn, when wheat residues are on the ground. The second parental strain may arrive (i) via pycnidiospores dispersed by rain splash or through direct contact from neighboring residues, or (ii) via ascospores carried by wind from more distant residues, as observed in 2019 and 2020 by Orellana et al. (2022). If we consider that sexual reproduction may already occur during the wheat growing period on senescent lower leaves as shown by Eriksen & Munk (2003) and Suffert et al. (2018), it is also conceivable that a pycnidiospore could land on a senescing leaf in late spring and cross with a virulent strain that had previously infected it. Still, this possibility can be reasonably considered epidemiologically negligible, given that the two strains involved would likely have had an earlier and more favorable opportunity to cross on living leaves, following splash-dispersal.

Scenario of type 4 corresponds to a variant of the PR treatment, in which the first parent is avirulent and does not infect the plant while it is alive. In this case, “one parent (avirulent strain) grows asymptomatically, possibly epiphytically, whereas the other parent (virulent or avirulent strain) arrives later on residues from a neighbouring set of residues” (Orellana et al., 2022). In the present study, a cross was obtained under such conditions; however, the suspected exogenous origin of the parental strains casts doubt on the relevance of this scenario, which, notably, had not been envisioned by Orellana et al. (2022). Interestingly, it aligns with the RR treatment — where both parents arrive later on residues — a case in which no offspring were obtained. This further questions the feasibility of such a reproductive pathway under natural conditions.

### 4.6. Final considerations

The picture we can form of what happens in the field is that of an equilibrium between various crossing scenarios, driven by processes — and therefore encounter conditions — that differ. The relative importance of each depends on agricultural practices, including physical contact and cross-contamination between varieties, which can be modulated in cultivar mixtures, as well as on whether residues are left on the ground. We have shown that scenario of type 3, although possible, is very likely quantitatively negligible. However, this raises a more fundamental question regarding the fact that sex can apparently occur independently of “infection” *sensu stricto*. Discussing the limits of sexual reproduction conditions provides another angle to reconsider the hemibiotrophic nature of *Z. tritici*, which until now has been based mostly on physiological aspects of the infection process (Sánchez-Vallet et al., 2015). Are the small proportion of crosses following scenario of type 3 a remnant of an ancestral phase of a saprotrophic, non-pathogenic species, or do they instead represent the most advanced stage of evolution in a biotrophic pathogen that originated as an endophyte? While our study does not delve into the mechanistic physiology of sexual reproduction, it broadens the scope of inquiry by considering its occurrence under agronomic conditions. Both approaches are complementary and deserve concurrent investigation to advance understanding of this issue.

## Research data

The research data used for the analysis are available in Supplementary Tables S1 and S2.

## Author contributions

FS developed the main conceptual framework, led the project, designed the experiments, and conducted the data analysis with input from MD and SLP. FS, MD and SLP carried out the *in planta* experiments, while SLP and AC performed the genotyping. FS wrote the manuscript in consultation with all authors.

## Declaration of generative AI in scientific writing

During the preparation of this work the authors used ChatGPT to improve the readability and language of the manuscript. The authors thoroughly reviewed, edited, and approved the final text and take full responsibility for the content of the published article.

## Supplementary material

**Supplementary Figure S1.**
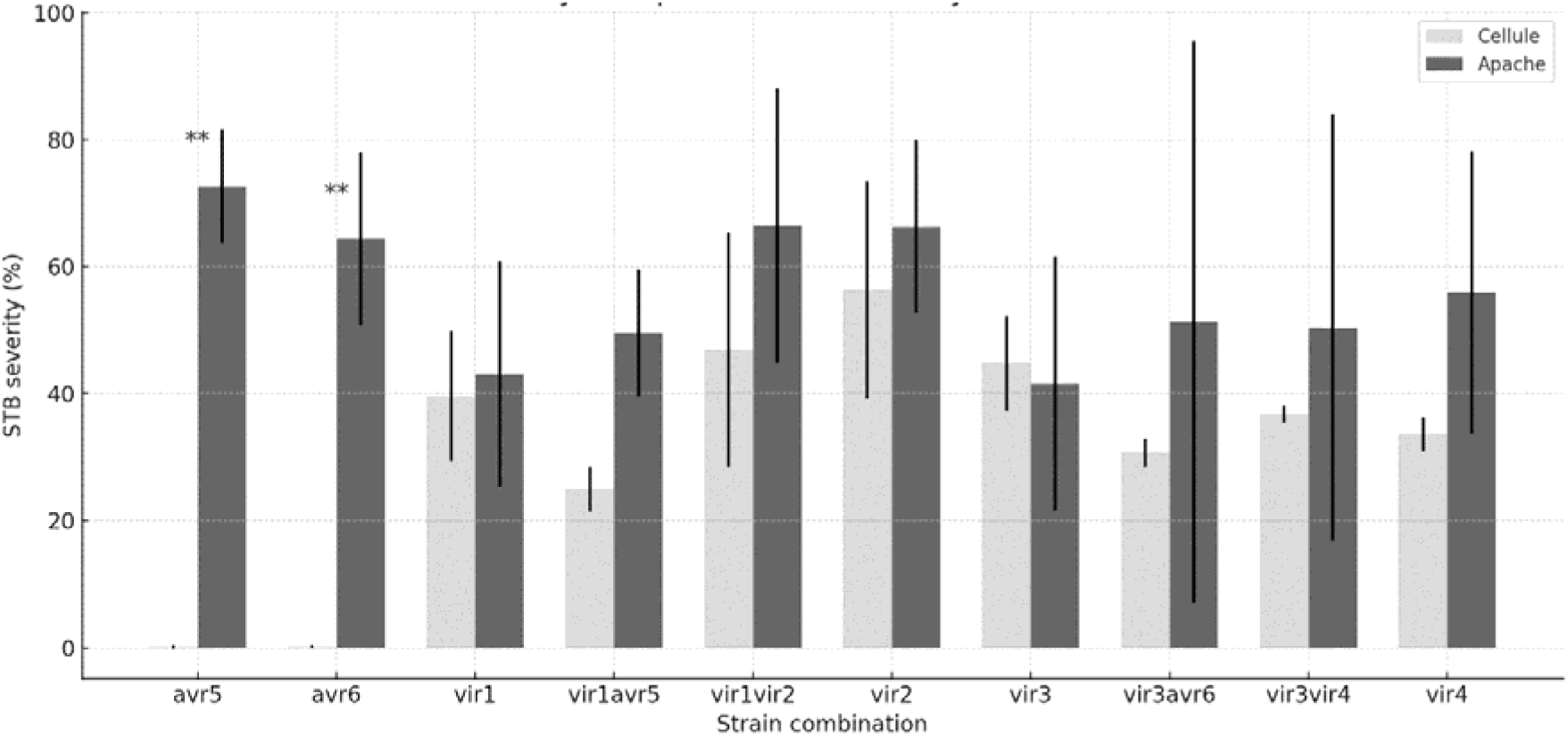
Comparison of STB severity levels across different strain combinations inoculated on living adult plant of Apache and Cellule. Mann–Whitney U test revealed significant differences only for single inoculations with avr5 and avr6 (p = 0.029).

